# A Degenerate PCNA-Interacting Peptide (DPIP) box targets RNF168 to replicating DNA to limit 53BP1 signaling

**DOI:** 10.1101/2021.03.17.435897

**Authors:** Yang Yang, Deepika Jayaprakash, Robert Hollingworth, Steve Chen, Amy E. Jablonski, Yanzhe Gao, Jay Ramanlal Anand, Elizabeth Mutter-Rottmayer, Jing An, Xing Cheng, Kenneth H. Pearce, Sophie-Anne Blanchet, Amélie Fradet-Turcotte, Grant S. Stewart, Cyrus Vaziri

## Abstract

The E3 ligase RNF168 has been suggested to have roles at DNA replication forks in addition to its canonical functions in DNA double-strand break (DSB) signaling. However, the precise role of RNF168 in DNA replication remains unclear. Here we demonstrate that RNF168 is recruited to DNA replication factories independent of the canonical DSB response pathway regulators and identify a degenerate PCNA-Interacting Peptide (DPIP) motif in the C-terminus of RNF168 which mediates its binding to PCNA. An RNF168 mutant harboring substitutions in the DPIP box fails to interact with PCNA and is not recruited to sites of DNA synthesis, yet fully retains its ability to promote DSB-induced 53BP1 foci. Surprisingly, the RNF168 DPIP mutant also retains the ability to support ongoing DNA replication fork movement, demonstrating that PCNA-binding is dispensable for normal S-phase functions. However, replisome-associated RNF168 functions to suppress the DSB-induced 53BP1 DNA damage response during S-phase. Moreover, we show that WT RNF168 can perform PCNA ubiquitylation independently of RAD18 and also synergizes with RAD18 to amplify PCNA ubiquitylation. Taken together, our results identify non-canonical functions of RNF168 at the replication fork and demonstrate new mechanisms of cross talk between the DNA damage and replication stress response pathways.

## Introduction

Ubiquitin signaling cascades play central roles in the sensing and repair of DNA damage. Failure to sustain DNA damage-inducible ubiquitin signaling compromises DNA repair resulting in increased sensitivity to genotoxic agents and genomic instability. DNA double-strand breaks (DSBs) such as those induced by ionizing radiation (IR) or radiomimetic agents are a potent inducer of the ubiquitin-dependent cellular DNA damage response (Ub-DDR), which is mediated primarily by the E3 ligases, RNF8 and RNF168. The mechanism of RNF8-RNF168 activation in response to DSBs is well-established (1-3) and is initiated by the DSB-responsive checkpoint kinase ATM which phosphorylates the histone H2A variant H2AX located in chromatin proximal to a break. Phosphorylated H2AX (termed γH2AX) recruits MDC1 which is also phosphorylated by ATM. The phosphorylation of MDC1 facilitates the recruitment of RNF8 to sites of damage, whereby it catalyzes the ubiquitylation of H1 and H2A-type histones surrounding the break. RNF8-dependent ubiquitylation of histones stimulates the recruitment of another E3 ligase complex, RNF168/UBC13 via direct interaction. Once recruited, RNF168/UBC13 mediates additional K63-linked poly-ubiquitylation of the ubiquitylated H2A histones. The sequential RNF8 and RNF168-dependent ubiquitylation of histones, together with other post-translational modifications already present at the break creates a platform for repair factors such as 53BP1, RNF169, and BRCA1 (4-6). 53BP1 is a critical DNA repair factor that dictates the pathway choice between NHEJ (non-homologous DNA end-joining) and homologous recombination (HR)-dependent DSB repair through its ability to suppress DNA end-resection, a process that is essential for initiating HR (7,8). The importance of the RNF168-dependent Ub-DDR in genome maintenance is evident from the existence of patients with ‘RIDDLE’ syndrome which is caused by bi-allelic mutations in *RNF168*. Patients with RIDDLE syndrome are immunodeficient and unable to produce IgG owing to defects in class switch recombination (CSR). Cells from RIDDLE syndrome patients are hypersensitive to ionizing radiation and fail to recruit 53BP1 to sites of DNA DSBs, recapitulating phenotypes of cells lacking H2AX, RNF8 and MDC1 (4,9).

In addition to the vital roles of ubiquitin signaling in the DSB response, ubiquitylation also plays key roles in sensing and processing of DNA damage or ‘stress’ that arises during DNA replication. DNA replication stalling can result from exogenously-induced bulky DNA lesions or DNA synthesis through highly repetitive regions of chromatin. The uncoupling of replicative DNA helicase and polymerase activities at stalled DNA replication forks generates ssDNA tracts (10) which are prone to breakage, a process often described as ‘fork collapse’.

To sustain S-phase and avert the lethal consequences of fork collapse, eukaryotic cells employ genome maintenance processes termed ‘DNA damage tolerance’ and ‘DNA damage avoidance’ which are also regulated by ubiquitin signaling cascades (11). The DNA damage tolerance pathway is triggered by activation of RAD18, an E3 ligase which interacts with ssDNA at stalled replication forks (12,13). RAD18 cooperates with the E2 ubiquitin conjugating enzyme RAD6 to mono-ubiquitylate PCNA at stalled replication forks (14). PCNA mono-ubiquitylation helps recruit specialized Y-family Trans-Lesion Synthesis (TLS) DNA polymerases to stalled replication forks (15). The Y-family DNA TLS polymerases (Polκ, Polη, Polι and REV1) have ubiquitin-binding domains and PCNA-interacting peptide motifs (termed ‘PIP boxes’) that favor PCNA binding when PCNA is in its ubiquitylated state (16). TLS polymerases are damage-tolerant when compared with the replicative DNA polymerases (Polδ, Polε) and are able to sustain DNA synthesis in cells harboring damaged genomes. TLS-deficient cells are intolerant of DNA damage and typically accumulate excessive ssDNA in S-phase which leads to checkpoint activation and lethal abortive mitoses (17,18). However, although TLS confers DNA damage tolerance, Y-family polymerases are error-prone and can increase the risk of mutagenesis. Each Y-family polymerase has a preferred cognate lesion that is bypassed with relative accuracy when compared with other TLS polymerases. For example, Polη bypasses UV-induced cyclobutane pyrimidine dimers (CPD) in an error-free manner and represses sunlight-induced mutagenesis (19). In Polη-deficient individuals (a congenital disease termed *xeroderma pigmentosum*-Variant or XPV), error-prone compensatory bypass of CPD adducts by alternative TLS polymerases leads to mutations and skin cancer-propensity (20).

Interestingly, RAD18-induced PCNA mono-ubiquitylation can also promote a relatively error-free DNA damage-avoidance pathway termed ‘Template Switching’ (TS) to sustain S-phase in cells with damaged genomes (21,22). TS uses fork-reversal as a mechanism for the leading strand to avoid damaged DNA and instead use the intact replicated lagging DNA strand as a template. In mammalian cells, the template-switching process requires two E3 ubiquitin ligases, HLTF and SHPRH, which further extend the K164-mono-ubiquitin moiety on PCNA, generating K63-linked poly-ubiquitin chains. K63-poly-ubiquitylated PCNA recruits helicases and chromatin-remodeling factors that mediate fork reversal and the template-switching process (23,24). Thus, ubiquitin signaling is crucial for normal responses to both DNA DSB and DNA replication stress.

Increasingly it is clear that E3 ubiquitin ligases are versatile enzymes that can ubiquitylate multiple substrates, have non-catalytic functions and participate in diverse biological processes. For example, RAD18 responds to DSB and potentiates HR via a non-catalytic mechanism involving chaperoning of the recombinase RAD51D to DSB sites (25). RAD18 was recently shown to mono-ubiquitylate histones in response to DSB (26). Although RAD18 is the main PCNA-directed E3 ubiquitin ligase in most cells, other E3 ligases including HLTF (27) and RNF8 may also mono-ubiquitylate PCNA (28). RNF168 was recently detected at DNA replication forks (29-31) and shown to promote normal S-phase progression (29). Therefore, E3 ligase-directed Ub-dependent signaling cascades are rarely specific to an individual type of DNA damage but rather form a complex network of integrated pathways that respond to a variety of different cellular stresses.

The mechanisms by which E3 ubiquitin ligases are targeted to different pathways potentially have enormous impact on DNA damage tolerance in cancer cells. Inappropriate expression levels of E3 ubiquitin ligases, or their inappropriate deployment to potential downstream substrates could lead to imbalance in DNA repair pathway choice that results in genetic instability and/or therapy-resistance. Interestingly, RNF168 and RAD18 are both aberrantly overexpressed in many cancers (32-35), although the impact of altered RNF168/RAD18 on tumorigenic phenotypes is poorly understood.

Given the impact of E3 ubiquitin ligases on DNA damage tolerance and genome stability in both normal development and pathological states it is important to understand how these enzymes are targeted to different genome maintenance pathways. Therefore, we sought to elucidate the mechanisms that target RNF168 to sites of DNA synthesis. Here we identify a degenerate PIP box (DPIP) located within the C-terminus of RNF168 that mediates its recruitment to replication forks. Using targeted mutagenesis, we demonstrate that the DSB and DNA replication functions of RNF168 are completely separable and that the DPIP motif is required to both limit the 53BP1-dependent DDR pathway at replication forks and promote ubiquitylation of PCNA at replication forks.

## Materials and Methods

### Cell culture and transfection

Cancer cell lines H1299, U2OS, 293T were purchased from the American Type Culture Collection (ATCC) and used for the described experiments without further authentication. U2OS *RNF168*^-/-^ and U2OS *RNF8*^*-/-*^ cell lines were gifted by Dr. Daniel Durocher, University of Toronto, Canada.

U2OS cells harboring Doxycycline-inducible human RNF168 were generated using the pINDUCER20 lentiviral vector (Addgene plasmid # 44012; http://n2t.net/addgene:44012; RRID:Addgene_44012). All cell lines were cultured in DMEM medium supplemented with 10% fetal bovine serum and penicillin–streptomycin (1%). All cell lines tested negative for mycoplasma contamination as determined using the ATCC Universal Mycoplasma Detection Kit (ATCC 301012K). Plasmid DNA and siRNA oligonucleotides were transfected using Lipofectamine 2000 (Invitrogen) according to the manufacturer’s instructions, except that plasmid DNA and Lipofectamine 2000 concentrations used in each transfection reaction were decreased by 50% to reduce toxicity.

### Adenovirus construction and infection

All other adenoviruses were constructed and purified as described previously (36). In brief, cDNAs encoding RNF168 variants were subcloned into the pACCMV shuttle vector. The resulting shuttle vectors were co-transfected with the pJM17 adenovirus plasmid into 293T cells.

Recombinant adenovirus clones were isolated by plaque purification and verified by restriction analysis and Southern blotting. The empty vector AdCon (used to control for adenovirus infections) was derived similarly but by co-transfection of the parental pACCMV shuttle vector with pJM17. Adenovirus particles were purified from 293T cell lysates by polyethylene glycol precipitation, CsCl gradient centrifugation, and gel filtration column chromatography. Adenovirus preparations were quantified by A260 measurements. Cells were typically infected with 0.01−3.0 × 10^10^ pfu/ml by direct addition of purified virus to the culture medium.

### RNA interference

siRNAs were delivered to cells using reverse-transfection or electroporation. For reverse-transfection, siRNAs were incubated with Lipofectamine 2000 and serum-free Optimem for 15 min at room temperature in the dark. Cells were then trypsinized, resuspended in 1 ml of Optimem and added directly into the siRNA/Optimem/Lipofectamine solution to give a plating density of 50%, then incubated for 48 hr. Electroporation was performed with a GenePulser Xcell (Bio-Rad), according to the manufacturer’s instructions. Typically electroporation was performed in a 0.2 cm cuvette containing 200 μl PBS, 2×10^6^ cells and 1μM siRNA using 150V (10 mS, 1 pulse).

siRNA sequences used here were: control non-targeting siRNA, 5’-UAGCGACUAAACACAUCAA-3’ (Thermo Fisher Scientific); RNF168 siRNA #1,siGENOME Smartpool Dharmacon (Cat# M-007152-03); RNF168 siRNA #2, 5’-GAGAAUAUGAAGAGGAAAUUU-3’; RNF168 siRNA #3, 5’-GAAGAGUCGUGCCUACUGAUU-3’ (described by Fradet-Turcotte et al., (5)); RAD18, 5′-GAGCAUGGAUUAUCUAUUCAAUU-3′; RNF8, 5′-GAGAAGCUUACAGAUGUUU-3′

### Immunoprecipitation and immunoblotting

To prepare extracts containing soluble and chromatin-associated proteins, monolayers of cultured cells typically in 10 cm plates were washed twice in ice-cold PBS and lysed in 500 μl of ice-cold cytoskeleton buffer (CSK buffer; 10 mM Pipes pH 6.8, 100 mM NaCl, 300 mM sucrose, 3 mM MgCl2, 1 mM EGTA, 1 mM dithiothreitol, 0.1 mM ATP, 1 mM Na3VO4, 10 mM NaF and 0.1% Triton X-100) freshly supplemented with Protease Inhibitor Cocktail and Phostop (Roche). Lysates were centrifuged at 4000 rpm for 4 min, to remove the CSK-insoluble nuclei. The detergent-insoluble nuclear fractions were washed once with 1 ml of CSK buffer and then resuspended in a minimal volume of CSK before analysis by SDS–PAGE and immunoblotting. For all immunoprecipitation experiments, input samples were sonicated and normalized for protein concentration. Magnetic beads containing covalently conjugated antibodies against the HA tag (M180-11, MBL) were added to the extracts and incubations were performed 30min at 4 °C using rotating racks. Immune complexes were recovered using magnetic stands. The beads were washed five times with 1 ml CSK (1 min per wash), to remove nonspecifically associated proteins. The washed immune complexes were boiled in protein loading buffer for 10 min, to release and denature for SDS– PAGE.

For immunoblotting, cell extracts or immunoprecipitates were separated by SDS-PAGE, transferred to nitrocellulose membranes, and incubated overnight with the following primary antibodies: PCNA (sc-56), RNF168 (polyclonal antibody raised against fusion proteins and affinity purified),^4^ β-actin (sc-130656), GAPDH (sc-32233), RNF8 (sc-134492) from Santa Cruz; pRPA32 S4/8 (A300-245A), Polη (A301-231A), Polι (A301-304A), Polκ (A301-977A) and RAD18 (A301-340A) from Bethyl Laboratories (Montgomery, TX); p-Chk1 S317 (2344), p-ATM S1891(4526) and H2A (2578) antibodies were purchased from Cell Signaling Technology; γH2AX (05-636) and H2A K15ub (MABE11119) were from Millipore; H2B (Ab1790) antibody was from Abcam. Antibody dilutions used for immunoblotting were 1:1,000, with exceptions for the following antibodies: PCNA (1:500), β-actin (1:5,000), GAPDH (1:5,000) and γH2AX (1:10,000).

### Live-cell Imaging and analysis

U2OS cells with mCherry-PCNA and Venus-53BP1 reporters were electroporated with siRNF168. RNF168-depleted cells were then seeded into 24-well glass-bottom plates (Cellvis, P24-1.5H-N) and 24 h later cells were infected with 1 × 10^8^ pfu/ml of Ad-RNF168 WT, Ad-RNF168 ΔDPIP or AdCon. Before imaging, cultures were switched to FluoroBrite(tm) DMEM (Invitrogen) media supplemented with 10% FBS, 4 mM L-glutamine, and penicillin/streptomycin. A high-content imager (InCell Analyzer 2200; Cytiva) was used to collect 12 fields of view from each well and the whole plate was imaged at 15-minute intervals for 20 hours with a 40x objective. The 4-megapixel CMOS sensor acquired the FITC and TexasRed channels set to a 1.5 s exposure and 2.5D imaging method set to 2 steps to deconvolute a 2-micron z-range around the target plane. Images were then analyzed and segmented using INCell Developer (Cytiva), cell tracking was performed using Visual Basic for Applications, and the resultant feature data was stored in a custom MySQL database.

### Immunofluorescence microscopy

24 h after adenoviral infection, plasmid transfection or doxycycline induction (20ng/ml) cultures were pulse labeled with Edu (Fisher #C10340) for 1 h, fixed with 2% PFA for 15 min, washed 3 times with PBS (5 min/wash), permeabilized with 0.2% Triton X-100 for 5 min, washed 3 times with PBS (5 min/wash), and blocked in PBS containing 3% BSA and 5% Donkey serum for 1 h. Next, the Edu click reaction was performed according to manufacturer’s instructions, followed by 3 5 min washes in PBS. Staining with the primary antibodies (1:200 dilution) was performed for 1 h. Then primary antibody solution was removed by washing 3 times in PBS (5 min/wash). Secondary antibody staining (1:300 dilution) was performed for 1 h. Secondary antibody was removed using 3 washes in PBS (5 min/wash). Finally Vectashield with DAPI (#H-1200 Vector laboratories) was applied to the stained cells prior to imaging. For experiments in which cells were stained with PCNA antibody, cells were fixed and permeabilized with ice-cold methanol for 20 minutes prior to antibody staining. For PLA assay, the cells were fixed and stained according to manufacturer’s instructions (Sigma Aldrich DUO92002). High resolution images were acquired by Zeiss LSM 710 Spectral or LSM 700 Confocal Laser Scanning Microscope at UNC Microscopy Service Lab (MSL) with 40x/60x objective. 100 nuclei per condition were counted manually for quantification.

### DNA fiber analysis

Sub-confluent cells were labelled with 25 µM chloro-deoxyuridine (CldU) (Sigma Aldrich) before media was removed and replaced with fresh media containing 250 µM iodo-deoxyuridine (IdU) (Sigma Aldrich). Cells were harvested at the indicated time, washed in PBS, added to glass slides and lysed in spreading buffer (200 mM Tris-HCl, pH 7.5, 50 mM EDTA, 0.5% SDS). DNA fibers were spread across the slides by tilting at an angle following lysis. Slides were air dried before fixation in methanol and acetic acid at a 3:1 ratio for 10 min. Prior to immunostaining, DNA was denatured using 2.5 M HCL for 1 h, 15 min before addition of blocking solution (1 % BSA, 1 % Tween 20 in PBS) for 30 min. Incorporated CldU and BrdU were detected using mouse anti-BrdU (clone B44; BD Biosciences, 347583) and rat anti-BrdU (clone BU1/75, ICR1; Abcam, ab6326) primary antibodies for 1.5 h. Slides were fixed with 4% paraformaldehyde for 10 min before addition of anti-rat Alexa Fluor 555 and anti-mouse Alexa Fluor 488 secondary antibodies (Thermo Fisher Scientific). Fluorescently-labelled DNA fibers were imaged using a Nikon Eclipse Ni microscope with NIS-Elements software (Nikon Instruments). Fiber track lengths were measured and recorded using ImageJ software (US National Institutes of Health; NIH). A minimum of 250 fibers were counted for each of three independent experiments.

### Isothermal titration calorimetry

Isothermal titration calorimetry (ITC; GE AutoITC 200) was conducted by titrating synthetic peptide (p21, GRKRR<u>QTSMTDFY</u>HSKRRLIFS-amide where underlined residues denote PIP box; syringe, 150 - 300 mM) into a solution of PCNA (cell; 30 mM) in TBS. Control experiments using peptide injected into buffer alone showed minimal heats with no evidence of titration. Analysis of the isotherm yielded Kd = 76.3 nM and stoichiometry = 0.93 (Microcal Origin software).

### In-vitro ubiquitylation assay

Ubiquitylation assays were performed in 25 mL reactions in which the components were added in the following order: dd H2O, 1x Energy regeneration solution (#B-10 R&D systems), 75 mM ubiquitin (#U-100H R&D systems) 1.6 mM FLAG-PCNA substrate (expressed and purified in bacteria), 0.1 mM E1 Ubiquitin Activating Enzyme (#E-304 R&D systems), 0.2 mM E2 conjugase (UbcH5c, #E2-627 R&D systems, or RAD6 #E2-613 R&D Systems), 0.2 mM E3 ligase (recombinant bacterial RAD18-RAD6 complex or RNF168 both purified in-house). Reaction mixtures were incubated at 37°C for 15 or 30 minutes, then inactivated by adding 1x Laemelli buffer and boiling for 10 minutes. Reaction products were then resolved on SDS-PAGE and analyzed by immunoblotting.

## Results

### RNF168 is recruited to DNA replication factories and DSB via independent mechanisms

To elucidate mechanisms of RNF168 recruitment to replication forks, we first asked whether the requirements for RNF168 localization to DSBs and replication factories were separable. It has been shown that the sustained accumulation of RNF168 at IRIF requires both the UMI and MIU2 ubiquitin binding domains (4).

Therefore, we compared the subcellular localization of WT RNF168 and a mutant lacking its MIU2 domain (ΔMIU2) in the absence of exogenous DNA damage and following exposure to hydroxyurea (HU) or ionizing radiation (IR). Consistent with previous observations (29), WT RNF168 co-localized with PCNA and this co-localization was enhanced in cells treated with HU but not IR (Fig. 1A, B). Interestingly, the RNF168 ΔMIU2 mutant which is compromised for recruitment to IRIF (4) displayed enhanced colocalization with PCNA when compared to the WT protein (Fig. 1A, B). Therefore, the mechanisms that recruit RNF168 to DSBs and sites of DNA replication stress are separable.

**Fig 1.**
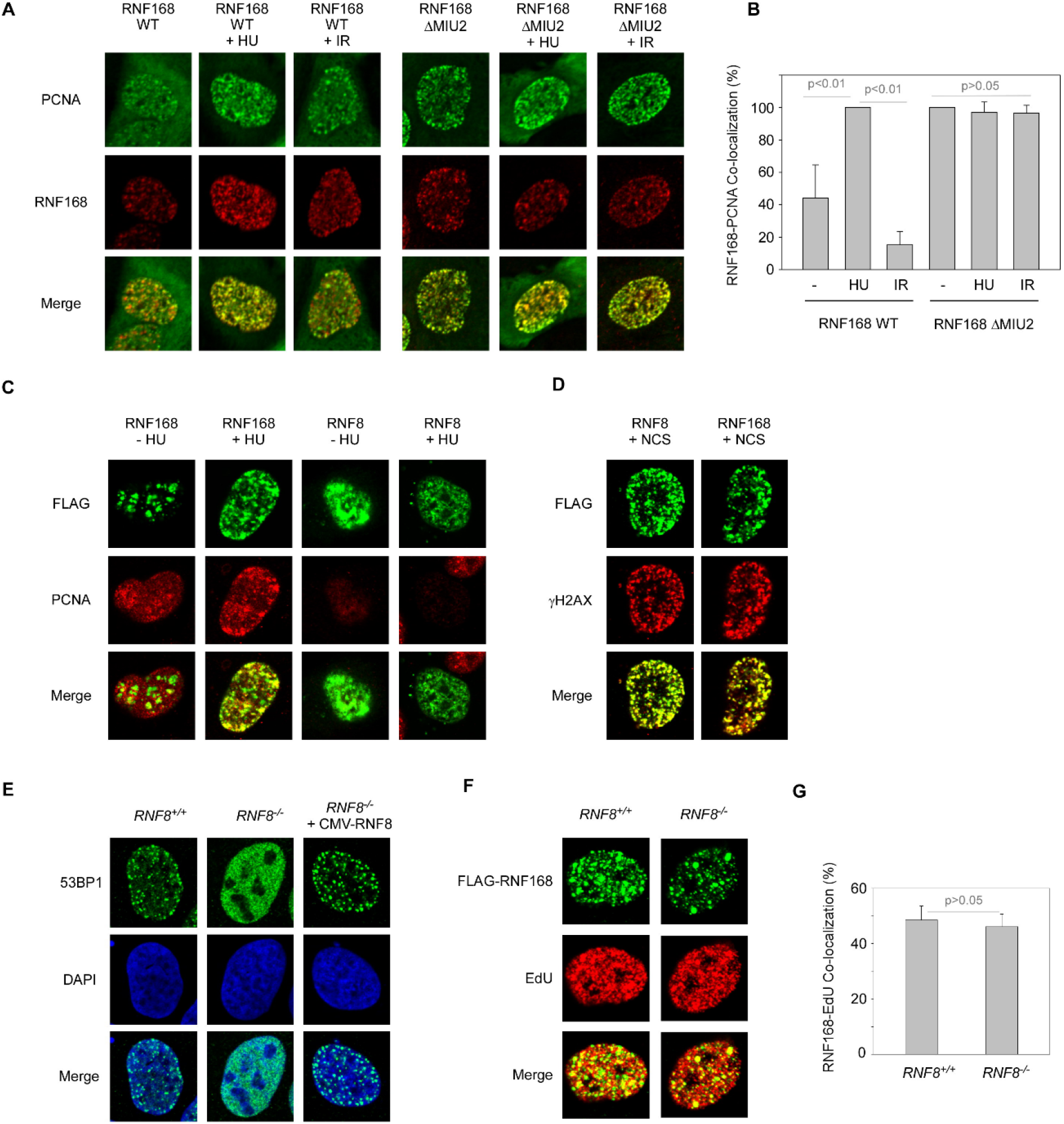
RNF168 is recruited to DNA replication and DSB via separable mechanisms. **(A)** U2OS cells were transiently co-transfected with expression constructs encoding HA-PCNA and FLAG-RNF168 WT or FLAG-RNF168 ΔMIU2. 48 h post-transfection cells were treated conditionally with 2 mM Hydroxyurea (HU) or with 10 Gy Ionizing Radiation (IR). Cells were fixed 2 h following HU or 30 min following IR treatments, then stained with anti-HA and anti-FLAG antibodies prior to analysis by immunofluorescence confocal microscopy. Panel (A) shows images of representative HA-PCNA and FLAG-RNF168 co-expressing cells from all experimental conditions. **(B)** Bar chart showing enumeration of cells containing HA-PCNA-co-localizing FLAG-RNF168 foci. Each data point represents the mean of results from 3 separate experiments and the error-bars represent the standard deviation. **(C)** U2OS cells were transiently co-transfected with constructs encoding FLAG-RNF168 WT or FLAG-RNF8 WT. 24 h post-transfection cells were treated conditionally with 2 mM Hydroxyurea (HU). Cells were fixed 2 h following HU treatment, then stained with anti-HA and anti-PCNA antibodies prior to analysis by immunofluorescence confocal microscopy. The individual images are of representative cells. Cells transfected with FLAG-RNF168 plasmids co-expressed both PCNA and RNF168 proteins. Unexpectedly however, in cultures co-transfected with FLAG-RNF8 plasmid, immunoreactivity with anti-PCNA and anti-FLAG antibodies was mutually exclusive (suggesting that FLAG-RNF8 is not expressed at detectable levels in PCNA-positive S-phase cells) **(D)** U2OS cells were transiently transfected with constructs encoding FLAG-RNF168 WT or FLAG-RNF8 WT. 24 h post-transfection cells were treated conditionally with 300 ng/ml Neocarzinostatin (NCS). Cells were fixed 1 h following NCS treatment, then stained with anti-FLAG and anti-γH2AX antibodies prior to analysis by immunofluorescence confocal microscopy. The individual images are of representative cells showing that both RNF8 and RNF168 co-localize with γH2AX. **(E)** Parental *RNF8*^*+/+*^ U2OS cells, *RNF8*^*-/-*^ U2OS cells, or *RNF8*^*-/-*^ cells transiently transfected with an RNF8 expression plasmid (for 24 h) were treated with 300 ng/ml NCS. 1 h after NCS treatment, cells were fixed and stained with 53BP1 antibodies prior to analysis by immunofluorescence confocal microscopy. The individual images are of representative cells showing RNF8-dependency of 53BP1 focus formation. **(F)** *RNF8*^*+/+*^ cells or *RNF8*^*-/-*^ derivatives were transiently transfected with an RNF168 expression plasmid. 24 h post-transfection, cells were pulse-labeled with 10 μM/ml EdU for 1 h to label sites of ongoing DNA synthesis. Cells were fixed and stained with antibodies against EdU and FLAG prior to analysis by immunofluorescence confocal microscopy. The individual images are of representative cells showing RNF8-independent co-localization of RNF168 and EdU. **(G)** Bar chart showing enumeration of cells containing EdU-co-localizing FLAG-RNF168 foci in *RNF8*^*+/+*^ and *RNF8*^*-/-*^ cells. Each data point represents the mean of results from 3 separate experiments and the error-bars represent the standard deviation. Unpaired student t test demonstrated no statistically significant co-localization between RNF8 and EdU incorporation (p = 0.5867).

Although it is known that RNF8 functions to promote RNF168 to sites of DSBs (4), it has been suggested that RNF8 can ubiquitylate PCNA and is therefore expected to be present at replication forks (28). However, while both RNF8 and RNF168 readily colocalized with γH2AX following exposure to neocarzinostatin (NCS), only RNF168 was recruited to replication forks in the presence/absence of replication stress (Fig. 1C, D). Despite this lack of detectable localization of RNF8 to sites of DNA synthesis, we hypothesized that the ATM-MDC1-RNF8 DDR axis could still potentially function to facilitate the recruitment of RNF168 to replication forks. To investigate upstream determinants of RNF168 recruitment to forks, we monitored the cellular localization of RNF168 in cells lacking RNF8. Interestingly, *RNF8*^*-/-*^ cells still retained the ability to recruit RNF168 to replication forks. Therefore, the canonical Ub-dependent DDR pathway controlled by RNF8 and RNF168 is specific to DSB signaling but the relationship between RNF8 and RNF168 diverges during DNA synthesis.

### The RNF168 C-terminus contains a novel PCNA-interacting motif

We sought to define the new non-canonical mechanism of RNF168 recruitment to sites of DNA replication. It has been previously demonstrated that RNF168 can bind PCNA in cultured cells (29). However, it is not clear whether RNF168-PCNA interaction is direct or mediated through its association with other replisome components. PCNA-binding proteins often associate with PCNA via PCNA-interacting Peptide (PIP) domains sharing the consensus sequence Q-X-X-L/V/I/M-X-F-F, where X represents any amino acid (37). We noticed that the C-terminus of RNF168 contains a degenerate PIP-like sequence Q-K-S-V-F-Q-M-F (Fig. 2A). Q559-F566 - resides in a C-terminal region FQAAQRCTK that is highly disordered (Fig. 2A) and therefore likely to mediate protein-protein interactions. Since PIP boxes tend to be located in disordered regions of PCNA-binding proteins (38,39) we reasoned that the PIP-like sequence may mediate the localization of RNF168 to replication forks.

**Fig. 2.**
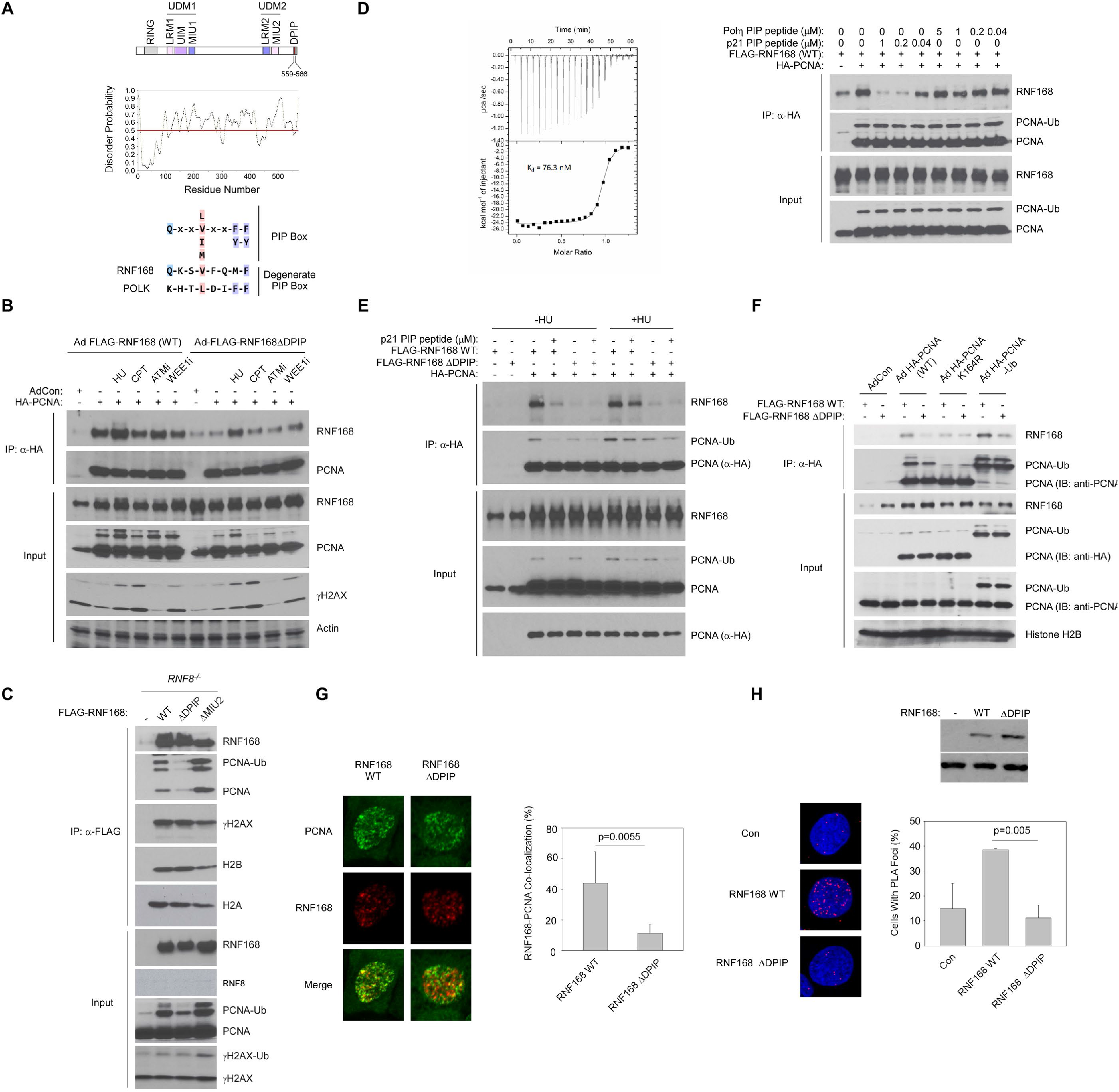
RNF168 contains a Degenerate PCNA-Interacting Peptide (DPIP) domain. **(A)** Domain organization of RNF168 indicating key functional domains and a degenerate PIP-like sequence residing in a disordered C-terminal region. The protein disorder profile was generated using the Protein Disorder Prediction (PrDOS) tool at http://prdos.hgc.jp/cgi-bin/top.cgi. **(B)** Replicate plates of H1299 cells were co-infected with adenovirus vectors encoding HA-PCNA and RNF168 WT or RNF168 ΔDPIP. 24 h post infection, some cultures were treated with HU (2mM, 2 h), camptothecin (CPT, 100 nM, 2 h), ATM inhibitor KU55933 100 nM, 2 h), or the WEE1 inhibitor MK1775 (10 μM, 2h). Chromatin extracts from the treated and untreated (control) cells were normalized for protein content and immunoprecipitated with anti-HA antibodies. Anti-HA immunoprecipitates were resolved on SDS-PAGE, transferred to nitrocellulose, and analyzed by immunoblotting with the indicated antibodies. **(C)** Replicate plates of *RNF8*^*-/-*^ U2OS cells were transiently infected with adenovirus vectors encoding FLAG-RNF168 WT, FLAG-RNF168 ΔDPIP, or FLAG-RNF168 ΔMIU2, or with a control ‘empty’ adenoviral vector. 48 h post-infection chromatin extracts were prepared, normalized for protein content and immunoprecipitated with anti-FLAG antibodies. Anti-FLAG immunoprecipitates were resolved on SDS-PAGE, transferred to nitrocellulose, and analyzed by immunoblotting with the indicated antibodies. **(D)** Isothermal titration calorimetry **(left panel)** was conducted by titrating synthetic peptide (p21, GRKRR<u>QTSMTDFY</u>HSKRRLIFS-amide where underlined residues denote PIP box; syringe, 150 - 300 μM) into a solution of PCNA (cell; 30 μM) in TBS. Control experiments using peptide injected into buffer alone showed minimal heats with no evidence of titration. Analysis of the isotherm yielded Kd = 76.3 nM and stoichiometry = 0.93 (Microcal Origin software). Replicate plates of H1299 cells were infected with adenovirus vectors encoding FLAG-RNF168 WT and HA-PCNA (or an ‘empty’ adenovirus vector for control). 36 h post-infection, chromatin extracts were prepared, normalized for protein content and immunoprecipitated with anti-FLAG antibody in the presence of different concentrations of peptides corresponding to the p21 or Polη PIP boxes. Anti-FLAG immunoprecipitates were resolved on SDS-PAGE, transferred to nitrocellulose, and analyzed by immunoblotting with the indicated antibodies **(right panel)**. **(E)** Replicate plates of H1299 cells were co-infected with adenovirus vectors encoding HA-PCNA and FLAG-RNF168 WT or HA-PCNA and RNF168 ΔDPIP. Some cells received an ‘empty’ adenovirus vector instead of HA-PCNA (for control). 36 h post-infection, some plates were treated with 2 mM HU for 2 h. Chromatin extracts were prepared, normalized for protein content and immunoprecipitated with anti-HA antibody in the presence or absence of the p21 PIP box peptide (1 mM). Anti-HA immune complexes were resolved on SDS-PAGE, transferred to nitrocellulose, and analyzed by immunoblotting with the indicated antibodies. **(F)** Replicate plates of H1299 cells were infected with adenovirus vectors encoding wild-type PCNA (HA-PCNA WT), ubiquitylation-resistant PCNA (HA-PCNA K164R), or a PCNA-ubiquitin fusion (HA-PCNA-Ub) in combination with FLAG-RNF168 WT adenovirus. Some cells received an ‘empty’ adenovirus vector instead of HA-PCNA viruses(for control). 36 h post-infection, chromatin extracts were prepared, normalized for protein content and immunoprecipitated with anti-HA antibody. Anti-HA immune complexes were resolved on SDS-PAGE, transferred to nitrocellulose, and analyzed by immunoblotting with the indicated antibodies. **(G)** H1299 cells were transiently co-transfected with expression constructs encoding HA-PCNA and FLAG-RNF168 WT or FLAG-RNF168 ΔDPIP. 48 h post-transfection cells were fixed, then stained with anti-HA and anti-FLAG antibodies prior to analysis by immunofluorescence confocal microscopy. The photographs are of representative cells co-expressing HA-PCNA and WT or ΔDPIP forms of FLAG-RNF168. The bar chart shows enumeration of cells containing HA-PCNA-co-localizing FLAG-RNF168 foci. Each data point represents the mean of results from 3 separate experiments and the error-bars represent the standard deviation. **(H)** *RNF168*^*-/-*^ U2OS cells were infected with adenoviral vectors encoding FLAG-RNF168 WT or FLAG-RNF168 ΔDPIP. 24 h post-infection cells were fixed and subject to proximity ligation assays to compare proximities of WT and ΔDPIP RNF168 with endogenous PCNA. The photographs are of representative DAPI-stained nuclei from each experimental condition. The bar chart showing enumeration of cells containing PCNA / FLAG-RNF168 PLA foci. Each data point represents the mean of results from 3 separate experiments and the error-bars represent the standard deviation. Ordinary one-way ANOVA demonstrated statistically significant difference between RNF168 WT and PIP (p = 0.0049).

To test the functionality of the putative DPIP box in RNF168 we complemented *RNF168*^*-/-*^ U2OS cells with FLAG-tagged RNF168 WT or an RNF168 mutant in which M565 and F566 were substituted with di-Alanine (hereafter designated ‘RNF168 ΔDPIP’), then co-immunoprecipitated PCNA and RNF168 from the resulting cell extracts.

Consistent with our immunofluorescence microscopy experiments showing RNF168-PCNA co-localization, WT RNF168 co-immunoprecipitated with PCNA and this interaction was moderately enhanced following exposure to HU but not CPT (Fig. 2B). Interestingly, disruption of the RNF168 DPIP box compromised its interaction with PCNA in unperturbed cells and significantly reduced its binding following HU treatment (Fig. 2B). These data indicate that the DPIP box in RNF168 primarily mediates its interaction with PCNA in unstressed cells but that in the presence of replication stress, RNF168-PCNA interaction also relies on another domain/motif.

Importantly, co-immunoprecipitation of RNF168 with PCNA was unaffected by pharmacological ATM inhibition (Fig. 2B). Moreover, the association of RNF168 with PCNA was robust and readily detected in cells lacking RNF8 (Fig. 2C). PCNA-binding also was unaffected by loss of the RNF168 MIU2 domain. Therefore, essential regulatory proteins (ATM, RNF8) and domains (MIU2) required to elicit the Ub-dependent DDR in response to DSBs are dispensable during S-phase. Of note, we observed that the RNF168 ΔMIU2 mutant bound more strongly to PCNA than WT RNF168, suggesting that the balance between the cellular pools of RNF168 is a binary choice dictated by binding either ubiquitylated histone or PCNA.

We reasoned that if RNF168 binds to PCNA via a PIP-box-mediated mechanism, the interaction would be competitively disrupted by a PIP box-containing peptide. We synthesized small peptide corresponding to the 20 amino acids sequences spanning the p21 PIP box and used ITC to confirm its association with PCNA (Fig. 2D, left panel). As shown in the Fig. 2D (right panel) the p21 PIP box peptide inhibited the amount of PCNA-co-immunoprecipitating RNF168 in a dose-dependent manner. In similar experiments, a peptide corresponding to the Polh PIP box (which binds PCNA with low affinity, Kd ∼400 nM) (40) when compared with p21 (Kd ∼76 nM) did not disrupt PCNA-RNF168 interactions as effectively (Fig. 2D). We conclude that the C-terminus of RNF168 contains a functional PIP box that mediates interactions with PCNA.

Interestingly, we noticed that the RNF168-PCNA interaction (as determined by co-IP) in HU-treated cells was more refractory to inhibition by the p21 PIP box peptide than the basal interaction in unstressed cells (Fig. 2E). Because HU induces PCNA mono-ubiquitylation (Fig. 2E), we hypothesized that covalent modification of PCNA by ubiquitin may further stabilize the RNF168-PCNA complex. By analogy, the association of PCNA with PIP box-containing TLS polymerases is also facilitated by mono-ubiquitylation of PCNA on K164 (16). To test the contribution of PCNA K164 ubiquitylation to RNF168 binding we used co-IP to compare association of RNF168 with wild-type PCNA and a K164R mono-ubiquitylation-resistant PCNA mutant. As shown in Fig. 2F, RNF168 WT showed reduced binding to PCNA K164R when compared with PCNA WT (indicating that PCNA modification at K164 promotes RNF168 binding). We also performed co-IP experiments to determine association of RNF168 with a PCNA-ubiquitin fusion protein that mimics mono-ubiquitinated PCNA (16). As also shown in Fig. 2F, RNF168 WT showed increased binding to PCNA-ubiquitin fusion protein when compared with PCNA WT. We conclude that RNF168 associates with PCNA via a degenerate PIP-box and that PCNA K164 mono-ubiquitylation further stimulates this binding.

Last, we utilized immunofluorescence and proximity ligation assays (PLA) to ascertain whether loss of the DPIP compromised the ability of RNF168 to be recruited to replication forks. As shown in Fig. 2G and Fig. 2H, ablating the DPIP box significantly compromised the re-localization of RNF168 to sites of ongoing DNA synthesis as judged by both immunofluorescence and PLA. Taken together, the results of Fig. 2 indicate that RNF168 is a *bona fide* PCNA-interacting protein.

### RNF168 supports S-phase progression independently of PCNA-interaction

It has been shown that RNF168 is required to maintain efficient DNA replication in the absence of exogenous replication stress (29). It was suggested that this function of RNF168 requires not only its E3 ligase activity and ubiquitin binding capabilities but also key members of the canonical Ub-DDR including H2AX, ATM, RNF8 and 53BP1. In light of our finding that RNF168 is specifically targeted to sites of DNA replication via its DPIP box, we reasoned that loss of PCNA-binding activity could potentially compromise the ability of RNF168 to sustain DNA synthesis.

To investigate the role of PCNA-binding in RNF168-dependent replication fork movement, we reconstituted *RNF168*^*-/-*^ U2OS cells with either WT RNF168 or with a mutant lacking the DPIP box and measured DNA replication dynamics using DNA fiber analysis. For comparison, we also complemented *RNF168*-null cells with an MIU2 domain mutant of RNF168. Consistent with previous observations, complementation of *RNF168*-null cells with WT RNF168 but not a mutant lacking the MIU2 domain restored replication fork speed (Fig. 3A, 3B). Strikingly, the RNF168 ΔDPIP mutant fully corrected the reduced fork speed of *RNF168*^*-/-*^ cells, conferring fork velocities comparable to those of WT RNF168-complemented cells (Fig. 3A, 3B). Identical results were obtained when we ectopically expressed WT and mutant forms of RNF168 using either adenoviral vectors (Fig. 3A) or doxycycline-inducible lentiviral vectors (Fig. 3B). The results of Fig. 3A-B suggest that the association of RNF168 with PCNA is not required to sustain efficient DNA replication rates.

**Fig. 3.**
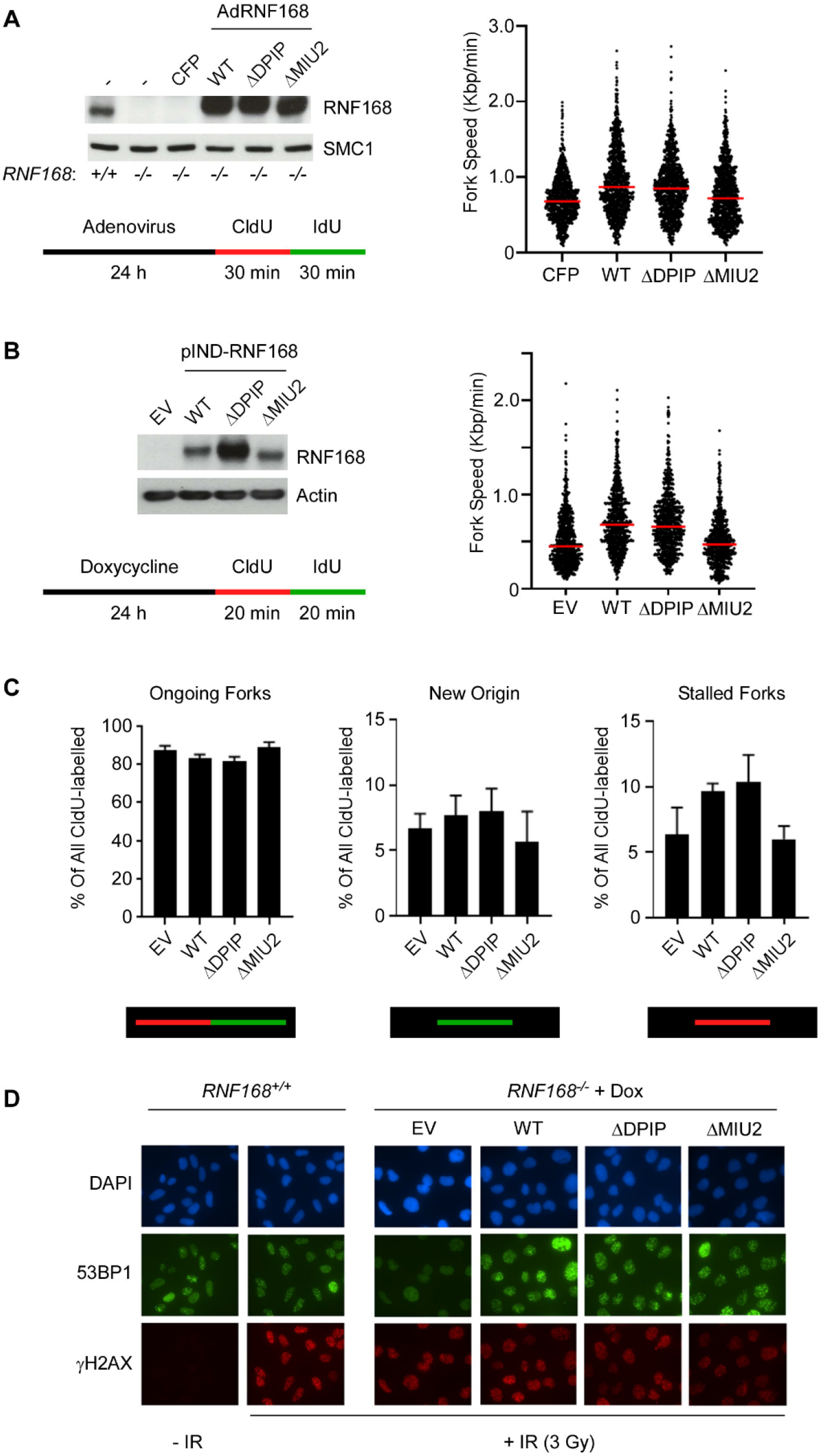
Wild-type and PCNA-interaction-deficient RNF168 sustain DNA replication. **(A)** Replicate plates of RNF168-deficient U2OS cells (*RNF168*^*-/-*^) were transiently transduced with adenoviral vectors encoding CFP (a control for infection efficiency), RNF168 WT, RNF168 ΔDPIP, and RNF168 ΔMIU2. 24 h after infection, one plate from each replicate pair was harvested for immunoblot analysis to verify complementation with WT and mutant forms of RNF168. The other replicate plates of cells were sequentially pulse-labelled with CldU and IdU. DNA fibers prepared from the labelled cells were immuno-stained with antibodies against CldU / IdU-treated and DNA replication dynamics were determined based on relative lengths of CldU/IdU-labelled tracts. **(B)** *RNF168*^*-/-*^ U2OS cells were stably transduced with retroviral vectors encoding doxycycline-inducible RNF168 WT, RNF168 ΔDPIP, and RNF168 ΔMIU2, or with an ‘empty’ retroviral vector for control. Replicate plates of cells were treated with 20 ng/ml doxycycline. 24 h after doxycycline treatment one plate from each replicate pair was harvested for immunoblot analysis to verify expression of WT and mutant forms of RNF168. The other replicate plates of cells were pulse-labelled with CldU and IdU and used to analyze DNA replication dynamics as described in **(A)** above. **(C)** Replicate cultures of cells from the same experiment described in panel (B) were stained with 53BP1 and γH2AX antibodies to confirm correction of the 53BP1 IRIF defect by the doxycycline-inducible forms of RNF168 in *RNF168*-null cells.

Given that 53BP1 has also been implicated in maintaining DNA synthesis under unperturbed conditions, we hypothesized whether the DPIP box of RNF168 may play a role in recruiting 53BP1 to replication factories. Interestingly, we did not observe any significant co-localization of 53BP1 with PCNA in the absence of exogenous replication stress irrespective of RNF168 status (Supplementary Fig. 1) suggesting that RNF168 does not play a significant role in recruiting 53BP1 to replication forks during normal DNA synthesis.

### RNF168-PCNA interactions limit DSB-induced 53BP1 foci during S-phase

It has been previously suggested that the replication-coupled dilution of the H4K20me2 epigenetic mark, which is essential for the binding of 53BP1 to chromatin, may function to limit the recruitment of 53BP1 to sites of DNA damage in S-phase, thus biasing the repair pathway choice towards an HR-dependent mechanism (31,41). Despite this, 53BP1 can still be recruited to DNA lesions in S-phase, albeit with a reduced capacity.

We reasoned that the PCNA-association of RNF168 during S-phase might provide a new mechanism for restraining RNF168 and limiting the recruitment of 53BP1 to DSBs. To test the role of RNF168-PCNA interactions in regulating 53BP1, we developed an accurate and quantitative live cell imaging platform for measuring RNF168-dependent and cell cycle phase-specific subcellular localization of 53BP1 in response to DNA DSB in asynchronous cultures.

An automated analysis pipeline was established to analyze images of U2OS cells co-expressing Venus-53BP1 and Cherry-PCNA reporters. Cell cycle phases were classified based on tracking mCherry-tagged PCNA. S-phase cells were identified based on distribution patterns of mCherry-tagged PCNA which forms characteristic nuclear foci during S-phase (Supplementary Figure 2) that are indistinguishable from S-phase-associated endogenous PCNA foci (42). Neocarzinostatin (NCS) was used to induce DSB and cells containing Venus-53BP1 foci were visualized and quantified. Fig. 4A shows quantification of 53BP1 distribution in representative NCS-treated U2OS cells.

**Fig. 4.**
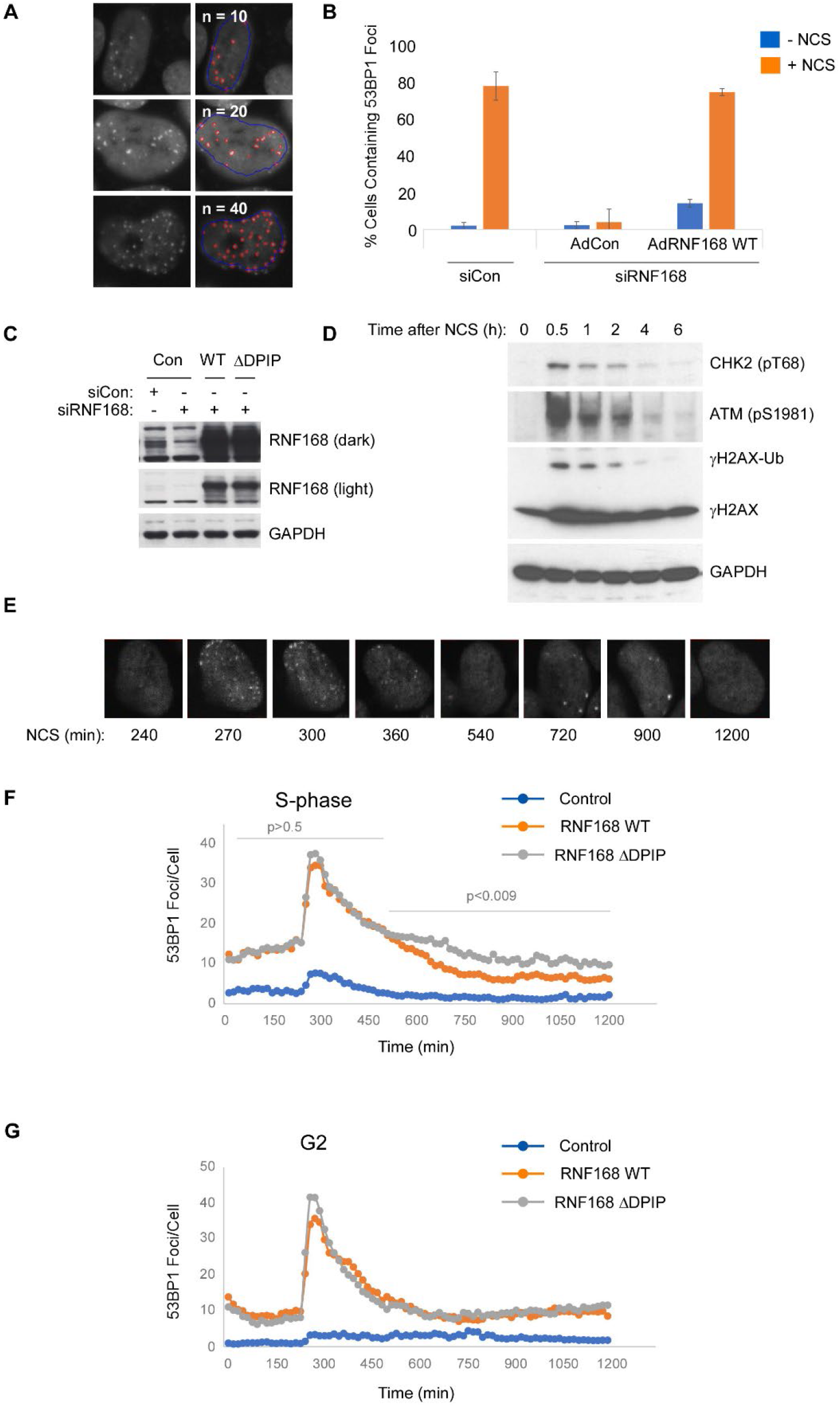
Wild-type and PCNA-interaction-deficient RNF168 support DSB-induced formation of 53BP1 foci. **(A)** Validation of automated image analysis pipeline for enumerating 53BP1 foci in representative NCS-treated cells. The three panels show segmentation results for three densities of 53BP1 foci. 53BP1 foci masks (red) were detected with granular segmentation and linked to the nuclear mask (blue). **(B)** U2OS cells containing mCherry-PCNA and Venus-53BP1 reporters were electroporated with siRNF168 or with non-targeting siCon RNA for controls. Control and RNF168-depleted cells were plated into imaging plate after electroporation and 24 h later cells were infected with 1 × 10^8^ pfu/ml of Ad-RNF168 WT or AdCon. 24 h post-infection, some cultures were treated with 100 ng/mL NCS and enumerated 53BP1 foci by confocal imaging. **(C)** U2OS cells containing mCherry-PCNA and Venus-53BP1 reporters were electroporated with siRNF168 or with non-targeting siCon RNA, re-plated after electroporation and infected 24 h later with 1 × 10^8^ pfu/ml of AdCon, Ad-RNF168 WT, or AdRNF168 ΔDPIP. 24 h post-infection cells were collected and analyzed using SDS-PAGE and immunoblotting with the indicated antibodies. **(D)** U2OS cells containing mCherry-PCNA and Venus-53BP1 reporters were treated with NCS (100 ng/ml) for the indicated times and analyzed using SDS-PAGE and immunoblotting for markers of the S-phase checkpoint. **(E)-(G)** U2OS cells containing mCherry-PCNA and Venus-53BP1 reporters were electroporated with siRNF168 and complemented with AdCon, Ad-RNF68 WT or AdRNF168 DPIP. Cultures were subject to live cell imaging every 15 min for 1200 minutes and were treated NCS (100 ng/ml) at 240 min. The panels in **(E)** show 53BP1 foci in a representative Ad-RNF68 WT-infected cell that was in S-phase at the time of NCS treatment and was repeatedly imaged for 1200 min. Panels **(F)** and **(G)** show mean numbers of 53BP1 foci from 3 independent experiments in which 50 cells in S-phase **(F)** or G2 **(G)** were individually tracked prior to and after NCS-treatment. In panel **(F)**, the mean numbers of foci per cell for Ad-RNF68 WT and AdRNF168 ΔDPIP-infected cultures are significantly different (p<0.05) at every time point after 480 min, as determined by 2-way repeated measures ANOVA with multiple comparisons.

Next, using the live cell imaging and automated analysis platform we established an assay for RNF168-dependency of NCS-induced 53BP1 focus formation. We used siRNA to deplete endogenous RNF168 from the Cherry-PCNA/Venus-53BP1 reporter cells and validated that RNF168-ablation led to loss of NCS-induced 53BP1 foci (Fig. 4B). We then identified the lowest expression level of adenovirally-expressed siRNA-resistant WT RNF168 protein that would fully correct the defective 53BP1 focus-formation activity in RNF168-depleted cells (Fig. 4B).

Having established a complementation assay for RNF168-dependent 53BP1 regulation, RNF168-depleted cultures were transduced with adenoviral vectors encoding equivalent levels of siRNA-resistant FLAG-RNF168 WT or FLAG-RNF168 ΔDPIP protein (Fig. 4C). The RNF168-depleted and RNF168 WT/ΔDPIP -reconstituted cells were then conditionally treated with NCS to induce DNA DSB. We used live-cell immunofluorescence microscopy to track and measure the dynamic NCS-induced redistribution of 53BP1 to foci in individual cells that were in S-phase at the time of drug treatment.

53BP1 foci were measured in individual NCS-treated cells every 15 minutes over 1200 minutes, a time period that spans the activation and the recovery phases of the DSB-induced S-phase checkpoint (evident from kinetics of NCS-induced ATM, CHK2 and γH2AX phosphorylation shown in Fig. 4D). Fig. 4E shows the dynamic changes in 53BP1 localization within a single representative (RNF168 WT-expressing) cell at different times after NCS treatment. As shown in Fig. 4F, the kinetics of NCS-induced 53BP1 focus formation, and the number of 53BP1 foci per cell were indistinguishable between RNF168 WT- and RNF168 ΔDPIP-expressing cells during the first 300 minutes of drug treatment. Interestingly however, approximately 240 minutes after NCS treatment, RNF168 ΔDPIP-expressing cells consistently maintained approximately twice as many 53BP1 foci/cell when compared with RNF168 WT-expressing cells. In contrast with cells that acquired DSB in S-phase, the presence of the ΔDPIP mutant RNF168 did not have any effect on the resolution of 53BP1 foci in cells that were in G2 at the time of NCS-treatment. These observations suggest that the RNF168-PCNA interaction serves to limit 53BP1 signaling specifically during the recovery phase of the DSB-inducible S-phase checkpoint.

The availability of cells co-expressing the Venus-53BP1 and mCherry-PCNA reporters also provided an opportunity to measure the dynamics and RNF168-dependency of 53BP1/PCNA co-localization. As shown in Supplementary Figure 3, cells in which 53BP1 and PCNA co-localized were relatively rare, both basally and after DNA damage. However, even in those rare cells, there was no significant difference in the extent or dynamics of 53BP1/PCNA co-localization when comparing RNF168 WT and RNF168 ΔDPIP-expressing cells. We conclude that the RNF168-PCNA interaction is dispensable for recruiting 53BP1 to PCNA in NCS-treated cells.

### RNF168-PCNA-interactions promote Trans-Lesion Synthesis (TLS)

To date, the only substrates reported to be ubiquitylated by RNF168 are H2A(X), BLM, 53BP1 and JMJD2A (43-46). Interestingly, in RNF168-overexpressing cells, we observed robust mono- and poly-ubiquitylation of PCNA even in the absence of exogenous replication stress and this ubiquitylation was dependent on the DPIP box (Fig. 5A).

**Fig. 5.**
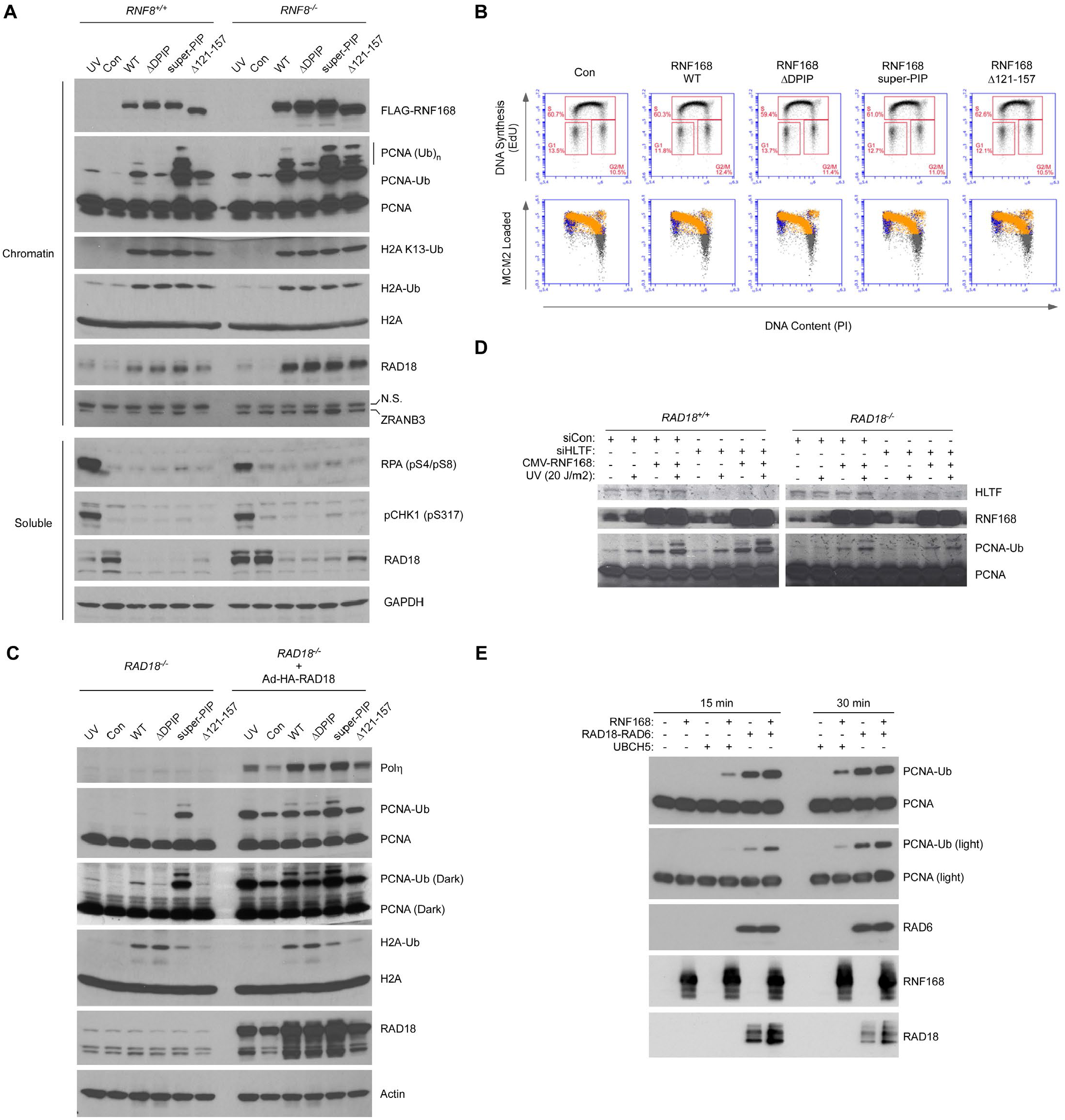
RNF168 synergizes with RAD18 to mono-ubiquitinate PCNA and stimulate TLS. **(A)** Replicate plates of *RNF8*^*+/+*^ and *RNF8*^*-/-*^ U2OS cells were infected with adenovirus vectors encoding RNF168 WT, RNF168 ΔDPIP, RNF168 super-PIP, and RNF168 Δ121-157, or with an empty adenovirus vector as control. After 23 h, one of the empty vector control plates was irradiated with UVC (60 J/m^2^). All plates of cells were collected 24 h post-infection and fractionated to give chromatin and soluble extracts. Cell extracts were normalized for protein content, resolved by SDS-PAGE and transferred to nitrocellulose membranes prior to immunoblotting with the indicated antibodies. **(B)** Replicate cultures of U2OS cells were infected with adenovirus vectors encoding RNF168 WT, RNF168 ΔDPIP, RNF168 super-PIP, and RNF168 Δ121-157, or with an empty adenovirus vector as control. After 23 h cells were labelled with EdU, extracted with nonionic detergent to remove unbound MCM, fixed, and stained with anti-MCM2 (a marker for the MCM2-7 complex), PI (total DNA), and for EdU incorporation (active DNA synthesis). Cell cycle phases are defined by DNA content (PI-A) and DNA synthesis (Edu-A) in the upper plots. Nuclei containing loaded MCM2 in G1 and S phase are represented by blue and orange dots respectively. G1/G2/M phase cells negative for chromatin-loaded MCM2 are shown in grey. **(C)** Replicate plates of *RAD18*^*+/+*^ and *RAD18*^*-/-*^ H1299 cells were infected with adenovirus vectors encoding different RNF168 variants (RNF168 WT, RNF168 ΔDPIP, RNF168 super-PIP, and RNF168 Δ121-157) in combination with RAD18 adenovirus, or with an empty adenovirus vector as control. After 22 h, two cultures were irradiated with UVC (60 J/m^2^). All plates of cells were collected 24 h post-infection and fractionated to give chromatin and soluble extracts. Cell extracts were normalized for protein content, resolved by SDS-PAGE and transferred to nitrocellulose membranes for immunoblotting with the indicated antibodies. **(D)** Replicate plates of *RAD18*^*+/+*^ and *RAD18*^*-/-*^ H1299 cells were sequentially transfected with HLTF-directed siRNA or with non-targeting control siRNA (siCon), then with CMV-FLAG RNF168 WT (or with an empty vector for control). 48 h post-transfection, some cultures were conditionally irradiated with UVC (20 J/m^2^). After 2 h chromatin fractions were prepared and analyzed by SDS-PAGE and immunoblotting with the indicated antibodies. **(E)** Purified PCNA substrate was incubated *in vitro* with recombinant RNF168 and recombinant UBCH5 individually or in combination, with recombinant RAD18-RAD6 complex, or with a combination of RAD18-RAD6 complex and RNF168 in the presence of E1, ubiquitin and an ATP-regenerating system. Reactions were terminated after 15 min or 30 min and products were separated on SDS-PAGE, transferred to nitrocellulose and analyzed by immunoblotting with the indicated antibodies.

To define the mechanism of RNF168-induced PCNA ubiquitylation, we expressed WT RNF168 and mutants lacking either the DPIP box or the UMI domain (Δ121-157) and monitored the ubiquitylation of H2A and PCNA. As a complementary approach to testing the effect of RNF168-PCNA association on PCNA ubiquitylation, we also expressed an RNF168 mutant in which the DPIP was replaced with the canonical high-affinity PIP box from p21 (QKSVFQMF>QTSMTDFY, hereafter termed ‘RNF168 super-PIP’). The WT and mutant forms of RNF168 were expressed in both *RNF8*^*+/+*^ and *RNF8*^*-/-*^ cells. Consistent with our previous observations, expression of WT RNF168 induced robust PCNA ubiquitylation that was ablated by inactivating mutations in the DPIP box. In contrast, mutating the DPIP to a high-affinity PIP box (super-PIP) significantly enhanced the RNF168-dependent ubiquitylation of PCNA. Therefore, the PCNA-binding ability of RNF168 is essential for RNF168-induced PCNA ubiquitylation. Strikingly, loss of the UMI domain of RNF168 (which prevents its binding to RNF8-ubiquitylated histone H1 and further recruitment of 53BP1 to DSBs) induced comparable levels of PCNA ubiquitylation to the super-PIP mutant in *RNF8*^*-/-*^ cells (Fig. 5A). The results of our mutational analyses (Fig. 5A) strengthen the idea that the DSB- and replication-associated functions of RNF168 exist in balance and disrupting one function enhances the other. Further indicative of a mutual competition between DSB- and DNA replication-related functions of RNF168, the ability of WT and ΔUMI RNF168 to stimulate PCNA ubiquitylation was enhanced in cells lacking RNF8. Interestingly, the RNF168-dependent ubiquitylation of H2A was unaffected by loss or upgrade of the DPIP or the presence/absence of RNF8, highlighting that the DPIP box is required to provide substrate specificity (i.e. favoring PCNA vs. H2A).

Since PCNA mono-ubiquitylation is a well characterized marker of replication stress, we investigated whether the increased ubiquitylation of PCNA following the expression of RNF168 arose indirectly through activation of the ATR-dependent replication stress response. However, in contrast to the robust phosphorylation of Chk1 and RPA2 in response to UV irradiation, Chk1 and RPA2 phosphorylation were not induced following the over-expression of WT or mutant RNF168 (Fig. 5A). However, we noticed that the expression of RNF168 was associated with the increased recruitment of RAD18 to chromatin (Fig. 5A). However, chromatin-binding of RAD18 did not correlate with levels of induced PCNA ubiquitylation, since the ΔDPIP, super-PIP and ΔUMI mutants of RNF168 all induced comparable levels of RAD18 recruitment (Fig. 5A) suggesting that perhaps the RNF168-induced increase in H2A ubiquitylation triggers RAD18 recruitment rather than promoting RAD18-mediated PCNA ubiquitylation.

To test the role of RAD18 in mediating RNF168-induced PCNA ubiquitylation we ectopically expressed WT and mutant forms of RNF168 in *RAD18*^*-/-*^ H1299 cells with or without co-expressed RAD18. As expected, the patterns of PCNA ubiquitylation induced by WT RNF168 relative to mutant RNF168 variants were similar in RAD18-replete cells (Fig. 5C, lanes 8-13) and *RNF8*^*+/+*^ and *RNF8*^*-/-*^ U2OS cells (Fig. 5A). Most unexpectedly however, RNF168-induced PCNA ubiquitylation was also readily detectable in *RAD18*^*-/-*^ cells (Fig. 5C lanes 1-6). Similar to results of experiments in RAD18-expressing cultures, PCNA ubiquitylation levels were lower in RNF168 ΔDPIP-expressing cells when compared with cells expressing RNF168 WT (and higher in RNF168 super-PIP-expressing cells when compared with cells expressing RNF168 WT). Although RAD18 co-expression clearly augmented RNF168-induced PCNA ubiquitylation (Fig. 5C), we conclude that RNF168 is capable of promoting PCNA ubiquitylation independently of RAD18. Although RAD18 is the predominant PCNA K164-directed E3 ligase, RAD18-mediated PCNA ubiquitylation can be augmented by the ligase activity of RNF168, in a manner that is governed by the affinity of its DPIP box.

The results of Fig. 5C suggested that RNF168 activates a PCNA-directed E3 ubiquitin ligase other than RAD18, or that PCNA is a direct target of RNF168-mediated E3 ubiquitin ligase activity. Other E3 ligases reported to ubiquitinate PCNA include RNF8 (28) and HLTF (27). Since RNF8 is dispensable for RNF168-induced PCNA ubiquitylation (Fig. 5A) we tested a possible relationship between HLTF and RNF168. However, siRNA-mediated depletion of HLTF did not affect RNF168-induced PCNA ubiquitylation in either WT or *RAD18*^*-/-*^ cells (Fig. 5D). Therefore, we performed *in vitro* ubiquitylation assays to test whether PCNA is a direct substrate of RNF168.

Given that RNF168 has been reported to form a complex with RAD6 (47), we initially used RAD6 as the E2 conjugating enzyme in the in vitro ubiquitylation reactions. However, we did not observe any significant ubiquitylation of PCNA by RNF168 in the presence of RAD6. Since RNF168 has also been shown to utilize UBCH5 as an E2 enzyme *in vitro*, we carried out the ubiquitylation reactions using UBCH5 instead of RAD6.

As shown in Fig. 5E, purified RNF168 readily mono-ubiquitinated ubiquitylated recombinant PCNA substrate in the presence of UBCH5 indicating that RNF168 can directly ubiquitylate PCNA. Interestingly, a combination of RAD18 and RNF168-UBCH5 led to a synergistic increase in PCNA mono-ubiquitylation *in vitro* (Fig. 5E), recapitulating the results obtained when we co-expressed RAD18 and RNF168 in cultured cells (Fig. 5C).

Taken together, the results of Fig. 5 identify RNF168 as a new mediator of PCNA ubiquitylation and potential activator of the TLS pathway.

## Discussion

Here we identify a new PCNA-interacting motif at the C-terminus of RNF168. We show that the PIP box is a novel functional domain that recruits RNF168 to sites of DNA synthesis and helps limit 53BP1 signaling during recovery from the DSB-induced S-phase checkpoint. Surprisingly, we also show that PCNA is a potential direct substrate of RNF168, and that RNF168-induced PCNA ubiquitylation in cells is dependent on the PIP box. In cells, RAD18 and RNF168 promote PCNA mono-ubiquitylation in a synergistic manner that can be recapitulated using the purified E3 ligases *in vitro*. What then is the basis for the cooperative PCNA ubiquitylation by RNF168 and RAD18? We hypothesize that RAD18 performs the initial ubiquitylation of one PCNA molecule within the PCNA trimer. Because RNF168 is preferentially recruited to ubiquitinated PCNA (Fig. 2), mono-ubiquitylation of one PCNA monomer is likely to stimulate association with RNF168 which then ubiquitinates other unmodified subunits of the PCNA trimer. Hu et al. previously showed that RNF168 and RAD18 both specifically bind histone H2A K13/15-ubiquitylated nucleosomes (48) and are therefore likely to coexist on chromatin. Because RAD18 and RNF168 both bind to ubiquitinated histones (and can both ubiquitinate histones), RNF168 and RAD18 signaling pathways may be intimately connected via both PCNA and histone modifications in a chromatin setting. For example, RNF168-PCNA interactions are facilitated by PCNA ubiquitylation (a modification that depends on RAD18). RNF168-dependent H2A-ubiquitylation in the vicinity of the replication fork could promote more RAD18 recruitment, potentially amplifying PCNA ubiquitylation (Fig. 6). Interestingly, Cipolla et al. show that Polη is recruited directly to ubiquitylated H2A. Because Polη is part of the RAD18 complex in cells, RAD18-associated Polη might also contribute to the recruitment of RAD18 to RNF168-modified chromatin.

**Fig. 6.**
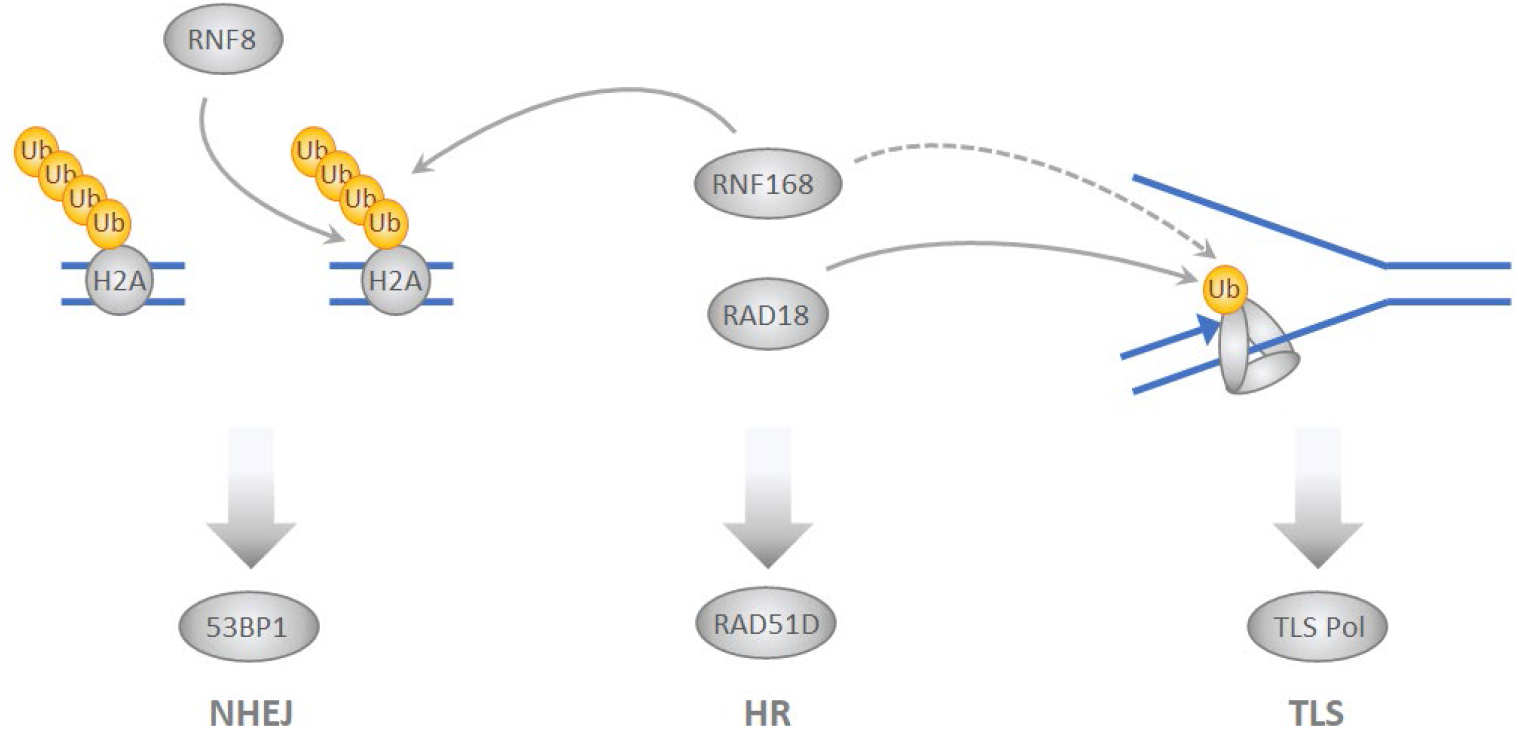
Summary of mechanisms by which RAD18 and RNF168 participate in genome maintenance at DSB (left) and DNA replication forks (right) RNF168 and RAD18 both ubiquitinate histone H2A in the vicinity of DSB to promote 53BP1 signaling and NHEJ **(left panel)** and also ubiquitinate PCNA to promote TLS **(right panel)**. RAD18 additionally acts as a molecular chaperone for the RAD51D recombinase and promotes HR independently of its ubiquitin ligase activity **(middle)**. See ‘Discussion’ for details.

The RNF168 and RAD18-mediated feed-forward loop that amplifies PCNA ubiquitylation is analogous to the cooperativity between RNF8 and RNF168 which serves to amplify histone ubiquitylation at DNA DSBs (1,2,49). Interestingly, PCNA ubiquitylation at DNA replication forks is also amplified via a cooperative interaction between RAD18 and Polη (50). Our results reveal yet another layer of intersection and crosstalk between the genome maintenance pathways that respond to DSB and DNA replication stress. For example, RAD18 was first described as a PCNA-directed E3 ligase that promotes TLS, yet was subsequently shown to promote DSB repair via HR (25,51), to facilitate chromatin retention of 53BP1 in G1 (52) and was recently identified as a histone-directed ubiquitin ligase (26). Conversely, RNF8 and RNF168 were first identified as mediators of the DSB response (4), yet both may also monoubiquitinate PCNA directly (see Zhang et al. (28) and this study) and thereby influence genome maintenance at the replication fork independently of DSB signaling. There are interesting parallels between RAD18- and RNF168-mediated ubiquitin signaling cascades and the ATM-CHK2 and ATR-CHK1 checkpoint kinase pathways; ATM-CHK2 and ATR-CHK1 were once considered to respond uniquely to DSB and replication stress respectively, yet it is now appreciated that these kinase cascades intersect and cross-talk extensively (53). Recent work (29,30,54) including this study demonstrates similar extensive crosstalk between DSB- and DNA replication stress-induced ubiquitylation cascades.

Having identified a new PCNA-dependent mechanism of RNF168 recruitment to sites of DNA synthesis, we sought to define the role(s) of RNF168 at DNA replication forks. Penengo and colleagues first showed that RNF168 is present at DNA replication forks in S-phase cells and proposed that RNF168 sustains normal DNA synthesis by recruiting 53BP1 and preventing excessive resection of reversed forks (29). Fully consistent with the results of Schmid et al. (29), we found that RNF168 confers normal rates of DNA replication fork movement. Surprisingly however, complementation experiments with our RNF168 PIP box mutant show that specific PCNA interaction is not required for RNF168 to promote normal replication fork movement. Consistent with the dispensability of RNF168-PCNA interactions for normal S-phase, Penengo and colleagues showed that the role of RNF168 in DNA replication was RNF8-dependent, whereas the interaction of RNF168 with PCNA is RNF8-independent. We surmise that global RNF168 activity may be sufficient for sustaining normal DNA replication fork rates, or that additional RNF8-dependent mechanisms exist that recruit RNF168 to replication forks.

Cipolla et al. also identified a connection between RNF168, PCNA ubiquitylation and TLS (30). Superficially similar to our results, Cipolla et al. showed that the elevated RNF168 activity in UBR5-depleted cells led to increased PCNA mono-ubiquitylation (30). However, those workers concluded that RNF168-induced PCNA ubiquitylation was a DNA damage response resulting from aberrant recruitment of Polη to DNA replication forks. In our studies, RNF168-induced PCNA ubiquitylation was clearly separable from the DNA damage response, and was not rescued by Polη-depletion.

Because RNF168 is a rate-limiting mediator of histone ubiquitylation following acquisition of DSB, there has been tremendous interest in determining the downstream targets and consequences of excessive RNF168 activity. RNF168 is overexpressed in neoplastic cells (34) including a subset of *BRCA1*-mutant cancers (35). Overabundance of RNF168 shifts DSB repair pathway choice from HR toward the relatively error-prone NHEJ (34). Remarkably, ectopically-expressed RNF168 promotes mutagenic NHEJ but does not affect physiological NHEJ-mediated repair of DSBs that arise during immunoglobulin class switch recombination or CSR (35). It was also shown recently that ectopic RNF168 promotes break-induced DNA replication in BRCA1-deficient cells (54). Clearly therefore, pathological changes in RNF168-mediated NHEJ are likely to contribute to the DNA damage tolerance and genetic instability of cancer cells (34,35).

Based upon our results showing that RNF168 stimulates PCNA mono-ubiquitylation, overactive RNF168 in cancer cells could confer not only chromosomal rearrangements and other forms of genetic instability that result from excessive NHEJ, but also increases in SNV burden due to aberrant activation of error-prone Y-family TLS polymerases. RAD18, the major PCNA-directed E3 ubiquitin ligase is also pathologically overexpressed and/or aberrantly activated in tumors (32,33) and levels of RAD18 mRNA correlate directly with overall SNV counts in many cancer types (33). It will be interesting to determine whether similar correlations exist between RNF168 expression and SNV burden in cancer. RNF168 and RAD18-induced mutability could be an enabling characteristic of cancer cells. Clearly further experiments are necessary to determine the impact of excessive RNF168 on mutability of cancer genomes.

In addition to ways in which pathological RNF168-PCNA interactions impact cancer biology, it is likely that RNF168-dependent PCNA-Ubiquitylation also participates in normal physiological genome maintenance. Raschle et al. performed mass spectrometry to study the dynamic recruitment of DNA repair factors to chromatin undergoing ICL repair (55). Those workers showed that Polη, RNF168 and RAD18 were coincident on chromatin during the TLS-mediated phases of ICL repair. Therefore it is tempting to speculate that the RAD18/RNF168 mediated amplification of PCNA ubiquitylation described here could influence the kinetics / efficiency of ICL repair.

PCNA mono-ubiquitylation is central to TLS polymerase activation, yet several proteins other than Y-family DNA polymerases associate with mono-ubiquitinated PCNA including Spartan (56) and SNM1A (57).

Moreover, the mono-ubiquitinated K164 residue of PCNA is further poly-ubiquitinated by human RAD5 homologues SHPRH and HLTF to promote template switching and DNA damage avoidance (27). It is probable therefore that RNF168-mediated PCNA ubiquitylation impacts not only TLS, but additional genome maintenance processes including spartan-mediated repair of DNA-protein crosslinks, SNM1-mediated ICL-repair, and template switching. Notably, the PCNA poly-ubiquitylation that mediates TS is a rare and unabundant post-translational modification that is typically challenging to detect using PCNA immunoblotting of whole chromatin extracts (27). In our experiments, RNF168 induced an unusually robust PCNA poly-ubiquitylation, perhaps hinting at a role for RNF168 in TS. Further experiments are underway to define the ways in which PCNA interactions influence RNF168 functions at the replication fork. The PCNA-interaction-deficient RNF168 DPIP allele described here is a very clean separation of function mutant that is intact for 53BP1 pathway regulation and will be a valuable tool for helping define additional contributions of RNF168-

PCNA signaling axis to DNA damage tolerance and genome maintenance.

## Supporting information

Supplemental figure RNF168

## Acknowledgements

We thank Dr. Natasha Zlatanou for suggesting the possibility that RNF168 associates with PCNA via a degenerate PIP box, Dr. Jeremy Purvis (University of North Carolina at Chapel Hill) for generous gift of mCherry-PCNA and Venus-53BP1 U2OS reporter cells, and Jitong Lou for data analysis suggestions.

## Supplementary Material

**Fig. S1** 53BP1 does not co-localize significantly with PCNA irrespective of RNF168 status and DNA damage.

**Fig. S2** Representative images of live-cell imaging experiments showing distribution patterns of mCherry-tagged PCNA throughout the cell cycle.

**Fig. S3** Frequency of co-localization of Venus-53BP1 and mCherry-PCNA in cells expressing RNF168 WT and ΔDPIP mutant forms.

