## Supplemental figure RNF168 for "A Degenerate PCNA-Interacting Peptide (DPIP) box targets RNF168 to replicating DNA to limit 53BP1 signaling"

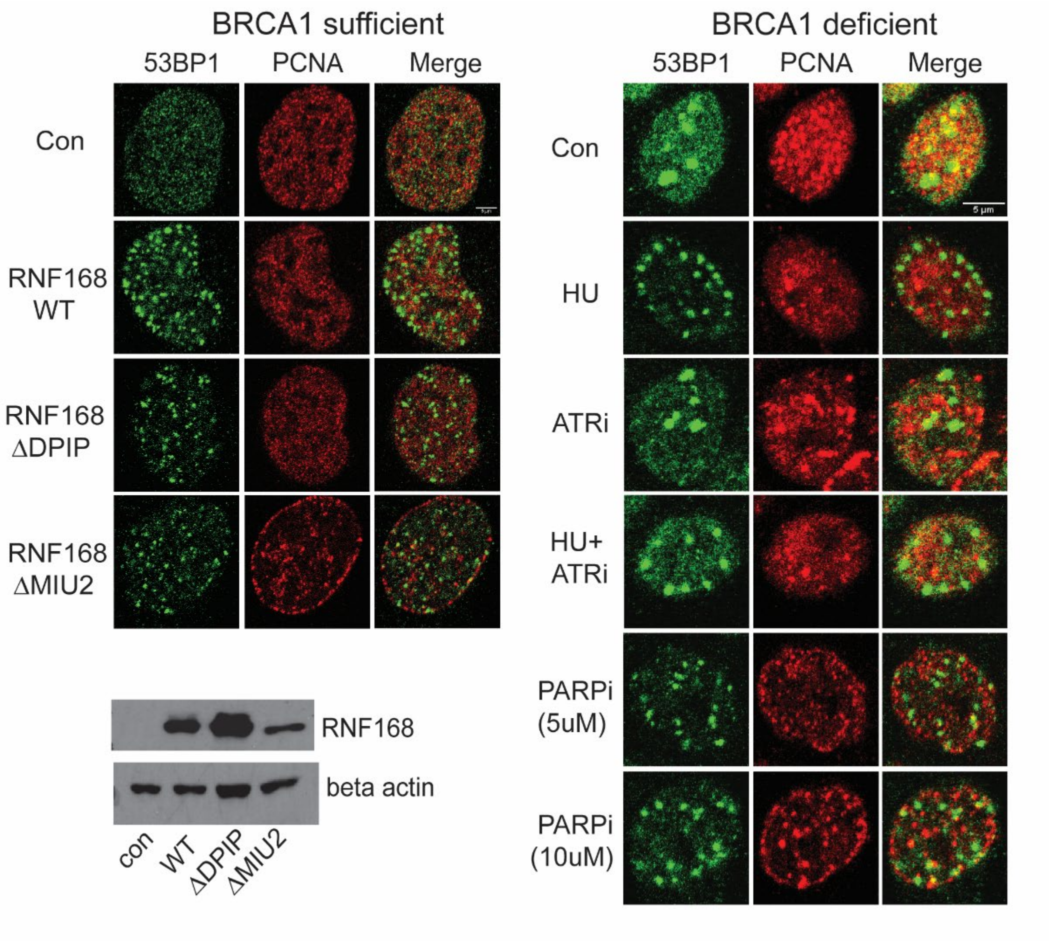


**Fig. S1** 53BP1 does not co-localize significantly with PCNA irrespective of RNF168 status and DNA damage.

<u>Left panel</u>: WT, ΔDPIP or ΔMIU2 RNF168 expression was induced with 20ng/ml dox in BRCA1 sufficient U20S RNF168KO pIND cells for 24 h. After 23 h, all conditions were subjected to 8 Gy ionizing radiation for 1 h before harvesting for immunoflourescnece to evaluate 53BP1-PCNA colocalization. A maximum of 4 foci/field of view at 40x showed colocalization of 53BP1 with PCNA irrespective of RNF168 status. The lower panel shows the relative protein expression of the RNF168 mutants by SDS-PAGE.

Right panel: Similar experiments were done in BRCA1 deficient MDA-MB-436 breast cancer cell line to evaluate if 53BP1-PCNA colocalization occurs as a function of BRCA1 status. The cells were infected with WT RNF168 adenovirus for 24 h and treated with the indicated DNA damaging agents for 2 h before harvesting for immunoflourescene (3mM HU, 3µM ATRi (inhibitor)). Irrespective of BRCA1 status and the type of DNA damaging agent used we again observed < 4 foci/field of view co-localization of 53BP1 with PCNA at 40x implying that 53BP1 does not significantly colocalize with PCNA foci.

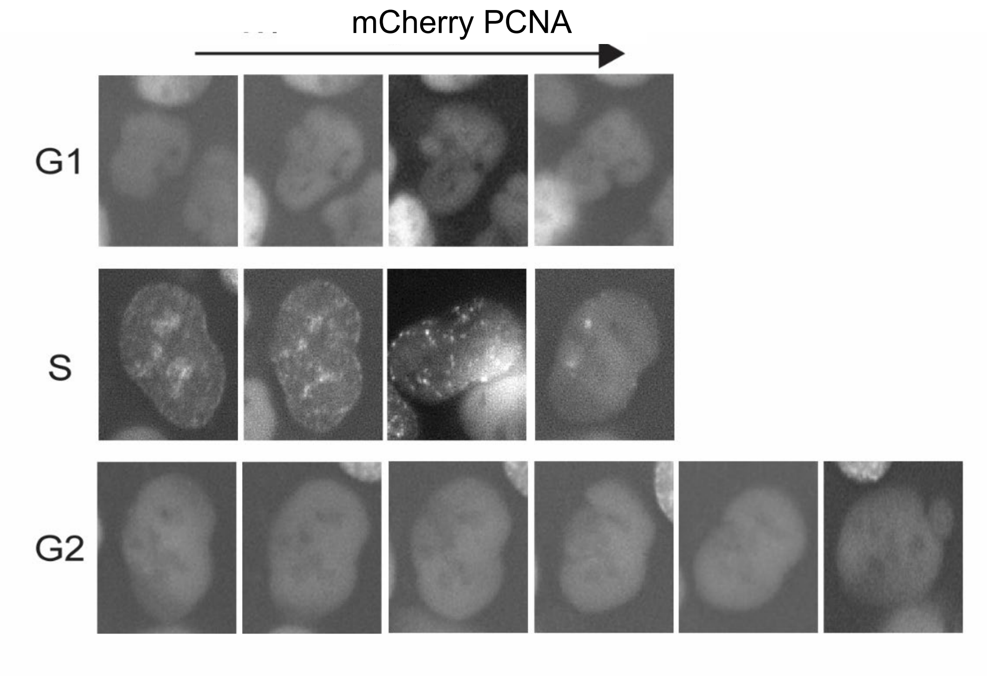


**Fig. S2** Representative images of live-cell imaging experiments showing distribution patterns of mCherry-tagged PCNA throughout the cell cycle.

The images represent the sequential progression of a single cell (U2OS reporter cell line) through G1, S and G2 phase as identified by the pattern of mCherry-PCNA foci expression in live cell imaging. The mCherry-PCNA foci pattern was used for segregating the tracked cells to different cell cycle phases through analytical pipeline generated (in INCell Developer software, Cytiva) for quantification of the experiment.

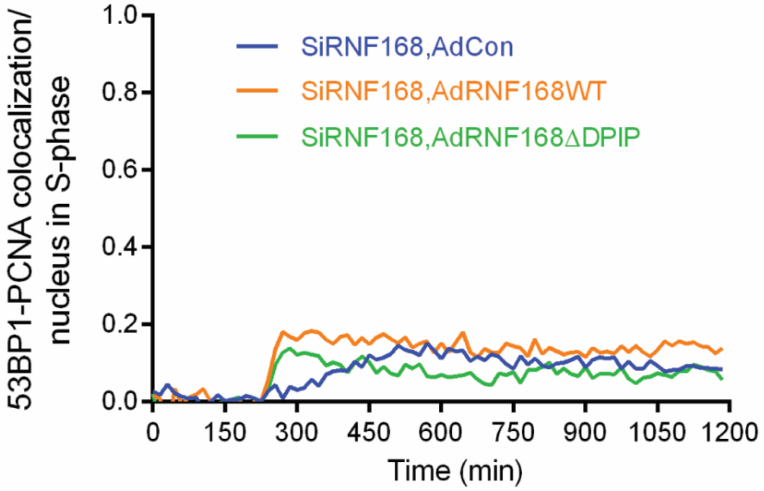


**Fig. S3** Frequency of co-localization of Venus-53BP1 and mCherry-PCNA in cells expressing RNF168 WT and ∆DPIP mutant forms.

U2OS cells containing mCherry-PCNA and Venus-53BP1 reporters were electroporated with siRNF168 and complemented with AdCon, Ad-RNF68 WT or AdRNF168 ΔDPIP. Cultures were then subjected to live cell imaging every 15 min for 1200 minutes and were treated with NCS (100 ng/ml) at 240 min. Mean number of 53BP1 and PCNA foci co-localization per nucleus in S-phase cells was automatedly analysed in INCell Developer using analysis pipeline demonstrating no significant co-localization.
